# SprayNPray: user-friendly taxonomic profiling of genome and metagenome contigs

**DOI:** 10.1101/2021.07.17.452725

**Authors:** Arkadiy I. Garber, Catherine R. Armbruster, Stella E. Lee, Vaughn S. Cooper, Jennifer M. Bomberger, Sean M. McAllister

## Abstract

Shotgun sequencing of cultured microbial isolates/individual eukaryotes (whole-genome sequencing) and microbial communities (metagenomics) has become commonplace in biology. Very often, sequenced samples encompass organisms spanning multiple domains of life, necessitating increasingly elaborate software for accurate taxonomic classification of assembled sequences. While many software tools for taxonomic classification exist, SprayNPray offers a quick and user-friendly, semiautomated approach, allowing users to separate contigs by taxonomy (and other metrics) of interest. Easy installation, usage, and intuitive output, which is amenable to visual inspection and/or further computational parsing, will reduce barriers for biologists beginning to analyze genomes and metagenomes. This approach can be used for broad-level overviews, preliminary analyses, or as a supplement to other taxonomic classification or binning software. SprayNPray profiles contigs using multiple metrics, including closest homologs from a user-specified reference database, gene density, read coverage, GC content, tetranucleotide frequency, and codon-usage bias. The output from this software is designed to allow users to spot-check metagenome-assembled genomes, identify, and remove contigs from putative contaminants in isolate assemblies, identify bacteria in eukaryotic assemblies (and vice-versa), and identify possible horizontal gene transfer events.

## 1. Background

There is demand among biologists for taxonomic classification and partitioning of genome and metagenome assemblies. In particular, there is a need for easy-to-use tools that allow novice users to more efficiently begin these analyses without sophisticated knowledge of programming languages like Python. Tools exist for taxonomic classification and tree-building, including those that use sets of specific gene markers on genomes or metagenome-assembled genomes (MAGs) binned from an original assembly (**Table 1**). The partitioning of contigs (i.e. binning) is often carried out independently of taxonomic-classification, and takes into account sequence compositional data (GC-content and tetranucleotide frequency) and read coverage; however, open-source software is available that can incorporate taxonomic/phylogenetic information to aid binning (Table 1). While many tools exist for classification of sequenced samples, several barriers exist for novice users, including the choice of which tools to use, how to use them, and the ability to efficiently iterate through these tools when analyzing multiple samples.

**Table 1.** Summary of published software for taxonomic classification and binning.

| Overall purpose | Software | Citation |
| --- | --- | --- |
| *Taxonomic classification and tree-building (specific markers)* | CheckM<br>PhyloSift<br>GToTree<br>phyloSkeleton | Parks et al., 2015<br>Darling et al., 2015<br>Lee, 2019<br>Guy et al., 2017 |
| *Taxonomic classification at the contig level* | Kaiju<br>Kraken<br>CLARK<br>FOCUS<br>MEGAN<br>BlobTools | Menzel et al., 2016<br>Wood and Salzberg et al., 2014<br>Ounit et al., 2015<br>Silva et al., 2014<br>Huson et al., 2016<br>Laetsch and Blaxter et al., 2017 |
| *Taxonomic classification at the contig level (all genes)* | CAT<br>Mmseqs2 | Meijenfeldt et al., 2019<br>Mirdita et al., 2020 |
| *Binning with sequence compositional data* | MetaBAT<br>MaxBin<br>Concoct<br>BinSanity<br>VizBin | Kang et al., 2019<br>Wu et al., 2014<br>Alneberg et al., 2014<br>Graham et al., 2017<br>Laczny et al., 2015 |
| *Binning with sequence compositional and taxonomic information* | Anvi’o<br>BlobTools | Eren et al., 2015<br>Laetsch and Blaxter et al., 2017 |

Here, we present an open-source bioinformatics tool, SprayNPray, that combines taxonomic/phylogenetic information with a variety of other metrics for each contig (discussed in detail below), allowing users to manually or automatically group contigs based on these metrics. This software wraps together several steps, which would otherwise require more advanced knowledge of a coding language to efficiently iterate through, and processes the output in a way that allows for easy visual inspection, manual curation, and/or further computational parsing. It is also organized in a way to allow for identification of pathogens, symbionts, and horizontally acquired genes in eukaryotic assemblies. The case studies presented below demonstrate the software’s versatility and potential usefulness to biologists dealing with non-axenic samples.

## 2. Implementation

SprayNPray, implemented in Python (version 3), is an easy-to-use software that provides a broad overview of input contigs by comparing each predicted ORF against a user-set reference database (recommended: NCBI’s non-redundant [nr] protein database [ftp://ftp.ncbi.nih.gov/blast/db/FASTA]). SprayNPray requires two inputs, a FASTA file of contigs (files with extension “.fna”, according to NCBI’s naming standards for genome FASTA sequences, but also .fa and .fasta in some cases) and a user-defined reference database, preferably NCBI’s non-redundant (nr) database of proteins. Users also have the option of providing a BAM file containing read coverage information; in this case, a script from the MetaBAT package (jgi_summarize_bam_contig_depths) is used to calculate average read depth per contig (Kang et al., 2019). After predicting ORFs with Prodigal (Hyatt et al., 2010), SprayNPray runs DIAMOND v2.0.4.142 (Buchfink et al., 2015) to query each ORF against the reference database (**Figure 1**). Results of this search are then grouped and written to a spreadsheet, where each row corresponds to a separate contig, followed by the taxonomic affiliation of the top DIAMOND hit to each ORF on that contig. SprayNPray ultimately writes three (optionally, four) output files:

1. In the main output file, SprayNPray provides the following metrics related to the supplied contigs:
  - Average amino acid identity (AAI) between the contig ORFs and closest matches in reference: this will provide users with an idea of how closely related their sequenced organisms are to what currently exists in public databases.
  - Number of genes normalized to the contig length: bacterial and archaeal genomes are typically gene-dense (1 gene per kbp), compared to eukaryotic genomes (0.9 genes per kbp in some fungi to 0.01 genes per kbp in plants and animals). Further, Prodigal is designed for prokaryotic ORF prediction, leading to suboptimal eukaryotic ORF prediction and subsequent lower gene density estimates. Thus, users can use coding density to deduce bacterial from eukaryotic contigs.
  - GC-content: if the provided contigs have organisms of varying levels of GC content, this will allow users to separate sequences based on that metric.
  - Read coverage: (this metric outputs only if a BAM file is provided): read coverage is useful for separating sequences if organisms represented among the contigs have varying levels of abundance, resulting in different read coverages estimates.
  - Contig length: this is a useful metric for filtering out low-quality contigs, or those too short to bin.
  - Cluster affiliation: contigs are clustered into putative bins. Cluster/bin assignments are derived from hierarchical clustering of contigs based on tetranucleotide frequency and codon usage bias. The number of clusters is estimated from the average number of hmmsearch (v.3.1b2, Johnson et al., 2010) hits per single-copy gene (using the ‘Universal_Hug_et_al’ set of 16 genes available in the GToTree package [Lee et al., 2019]).
2. SprayNPray provides an R-generated word cloud that is based on the distribution of the top taxonomic hits to genes predicted from the provided contigs (**Supplemental Figure 1**). When users provide a BAM file along with their contigs, the word sizes in the word cloud are corrected with read coverage information.
3. SprayNPray provides a file containing the top user-specified number of hits (default=100) to each ORF, allowing users to assess the taxonomic and functional distribution of top homologs to each gene.
4. SprayNPray has the capacity to write FASTA files that represent subsets of the provided contigs. This capability allows users to easily extract contigs belonging to organism(s) of interest (e.g. contaminants, pathogens, symbionts, certain genera). Subsets are created based on user-specified parameters, including: GC-content, coding density, amino acid identity, contig length, read coverage, and cluster/bin affiliation. Additionally, users can directly specify a taxonomic group of interest, and SprayNPray will write a FASTA file containing only contigs where some user-specified percentage (e.g. > 50%) of DIAMOND hits are to that specified taxa. In the event that the parameters by which FASTA files need to be written are unknown prior to running SprayNPray, users can re-run the program, with newly specified parameters (inferred from visually inspecting the output file from a preceding run), with greatly reduced runtime by providing the DIAMOND BLAST output file (file with extension .blast) from the previous run.
5. When running SprayNPray on an assembly containing eukaryotic contigs, users can also direct the program to specifically look for potential horizontal gene transfers (HGTs) from Bacteria or Archaea to Eukaryota. In this case, SprayNPray will write a separate output file containing putative HGTs. To identify ORFs of possible bacterial or archaeal origin, SprayNPray evaluates the taxonomic distribution of the top user-specified (default=100) DIAMOND matches for each ORF on each eukaryotic contig, and if more than a user-specified percentage (default 50%) of the hits are to bacterial proteins, that ORF is flagged as a potential HGT of bacterial or archaeal origin. In order for this part of the software to function properly, users need to be sure to include a reference database that encompasses protein sequences from all domains of life (e.g. nr).

**Figure 1:**
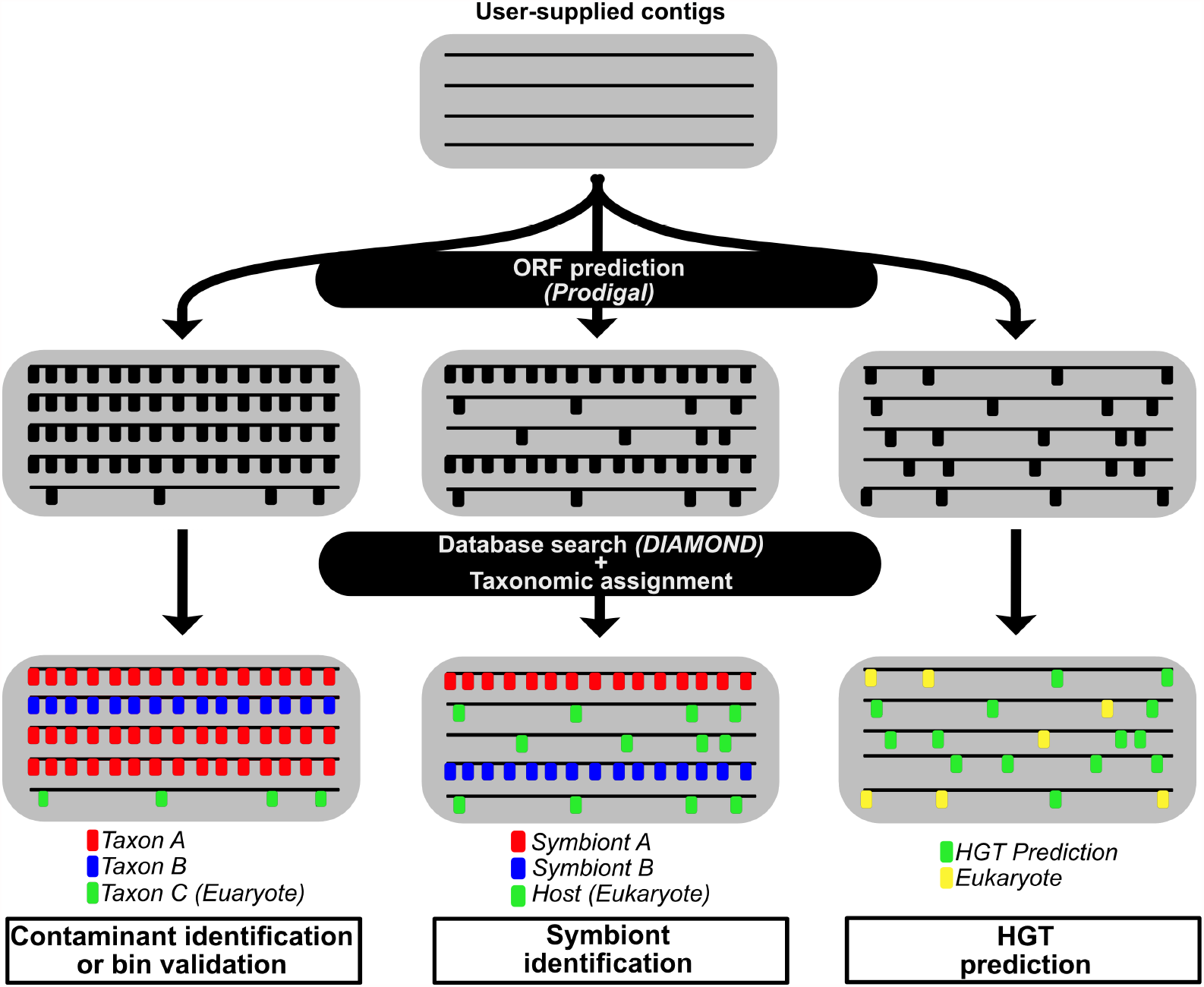
Overall workflow of the SprayNPray pipeline, with the four different uses (contaminant identification, bin validation, symbiont identification, and HGT prediction) shown. Horizontal lines in each gray box represent contigs, while the smaller vertical lines perpendicular to the contigs represent ORFs.

## 3. Case Studies

### 3.1 Bin validation

SprayNPray was used to profile a subset of metagenome bins from the North Pond aquifer (Tully et al., 2018) using NCBI’s nr database (release 200) as reference. To simulate the SprayNPray run as if it were conducted on an unpublished metagenome, we removed top hits corresponding to proteins generated from the original publication (Tully et al., 2018). The results (**Supplemental Files 1-4**) demonstrate SprayNPray’s capacity to 1) provide a rough taxonomic prediction of a genome bin and 2) demonstrate that the apparent taxonomic heterogeneity of a bin is due not to contaminating contigs, but to the low representation of a genome in NCBI’s nr database. This deduction is supported by the low average amino acid identity between the ORFs predicted in each contig and their closest hits in NCBI’s RefSeq database (**Supplemental Files 1-2**). Nonetheless, a subset of genome bins with higher similarities (>85%) to reference proteins recruited a more taxonomically homogenous set of homologs from the reference database (**Supplemental Files 3-4**), and appear less “contaminated”.

### 3.2 Contaminant identification in cultured isolates

SprayNPray can be used to identify putative contaminants in isolate assemblies. This software was used to remove contaminating *Serratia marcescens* contigs from a genome assembly of *Pseudomonas aeruginosa* isolated from a mixed clinical specimen containing multiple species. Visual inspection of the software’s output revealed the presence of contigs from both species (**Supplemental Files 5-7**). Running SprayNPray on these assemblies with the ‘-genus Pseudomonas’ flag created new FASTA files with only sequences corresponding to the isolate-of-interest, *Pseudomonas aeruginosa* (**Supplemental Files 8-10**), while contigs where >50% of hits were to homologs not affiliated with *Pseudomonas* were written to a separate file, which corresponded to the contaminating *Serratia* sequences (**Supplemental Files 11-13**). Subsequent analysis with CheckM v1.1.1 (Parks et al., 2015) confirmed the lack of contamination in the FASTA files written by SprayNPray (**Supplemental File 14**), while the completeness score for the newly written, clean *Pseudomonas* assemblies remained >99%, indicating that none of the *Pseudomonas*-sequences were removed.

### 3.3 Symbiont identification

SprayNPray can be used to extract bacterial symbiont contigs from an assembly that contains DNA from a variety of sources and domains of life. As an example, SprayNPray was run on an assembly of *Maconellicoccus hirsutus* (Kohli et al., 2020), the hibiscus mealybug, which contains two bacterial endosymbionts (Husnik and McCutcheon et al., 2016). In this assembly, the majority of DNA is from the host insect. Visual inspection of the initial output of this assembly (**Supplemental File 15**) allowed for the identification of metrics (e.g. GC-content, top taxonomic hits, gene density) with which the software was re-run to generate two additional FASTA files, each corresponding to an individual endosymbiont (**Supplemental Files 16-17**).

### 3.4 HGT identification

SprayNPray can be used to search eukaryotic contigs for genes that may have been horizontally/laterally obtained from bacteria via HGT. To showcase this functionality, we ran SprayNPray on an assembly of the citrus mealybug, *Planococcus citri* (de la Filia et al., 2021) (**Supplemental Files 18-19**), which is known to encode multiple HGTs from bacteria on its nuclear genome (Husnik et al., 2013). A total of 519 putative HGTs were identified (**Supplemental File 20**), including those previously confirmed by Husnik et al., 2013 (e.g. *murACDEF, bioABD, dapF, ddl*), as well as those that were reported but not confirmed (e.g. numerous AAA [ATPases Associated with diverse cellular Activities]-family ATPases of diverse origins, ankyrin repeat proteins with close homology to *Wolbachia* spp., and type III effectors) (Husnik et al., 2013). We note, however, that extreme caution should be taken in interpreting candidate HGTs, which should be validated by exploring the genomic context of each putative gene that is thought to have been horizontally acquired.

## 4. Conclusions

Here, we present SprayNPray, a bioinformatics software designed to aid in the taxonomic analysis of diverse (meta)genomic datasets. The appeal of this software is its ease-of-use and straightforward output that is amenable to visual inspection and/or computation parsing. We designed this versatile software to lower barriers for those with limited experience in bioinformatics and programming.

## 5. Materials and Methods

### Data acquisition

Metagenome-assembled genomes (MAGs) from the North Pond aquifer were obtained from the figshare link provided by Tully et al. (2018): (https://figshare.com/s/939160bb2d4156022558). Genomic data from the mealybug *Maconellicoccus hirsutus* were obtained from the following Genbank accession number: GCA_003261595.1 (Kohli et al., 2020). The genome assembly of the mealybug *Planococcus citri* were obtained from the MealyBugBase download server (https://download.mealybug.org).

### Isolation and sequencing of Pseudomonas aeruginosa

*P. aeruginosa* was isolated from clinical specimens following an IRB-approved protocol (STUDY19100149) at the University of Pittsburgh. Specimens were streaked onto *Pseudomonas* Isolation agar (PIA), a *Pseudomonas*-selective media on which *Serratia marcescens* and some other species can also grow, and incubated at 37oC for 48 hours. Single colonies were stored as a 30% glycerol stock at -80°C. Genomic DNA was extracted using a QIAgen DNeasy kit (Qiagen, Hilden, Germany) and sequenced on an Illumina NextSeq 500, with a target of 80-fold mean read coverage. Reads were trimmed with Trimmomatic version 0.36 1 in paired end mode, removing reads shorter than 70, quality checked with FastQC v0.11.5 (https://www.bioinformatics.babraham.ac.uk/projects/fastqc/). Trimmed reads were assembled using SPAdes v3.11.0 (Bankevich et al., 2012) and contigs smaller than 1kbp were removed.

## Availability and Requirements

**Project Name:** SprayNPray

**Project home page:** https://github.com/Arkadiy-Garber/SprayNPray

**Operating system(s):** Linux/MacOS

**Programming language:** Python

**Other requirements:** Python3, DIAMOND, Prodigal, MetaBAT

**License:** GNU General Public License v3.0

**Any restrictions to use by non-academics:** No further restrictions to use beyond license.

## List of Abbreviations

GC content: guanine cytosine content, represented as a percentage of total base pairs
MAGs: metagenome-assembled genomes
ORF: open reading frame
NCBI: National Center for Biotechnology Information
nr: NCBI’s non-redundant protein database
BAM: binary alignment map (file type)
AAI: average amino acid identity
kbp: kilobase-pair
BLAST: basic local alignment search tool
HGT: horizontal gene transfer

## Declarations

### Ethics approval and consent to participate

Not applicable

### Consent for publication

Not applicable

### Availability of data and materials

Supplemental files, corresponding to the genome data used and SprayNPray output, are provided with this article. These data are also freely available in the GitHub repository of this software: https://github.com/Arkadiy-Garber/SprayNPray/tree/master/Supplement-Case_Studies.

### Authors’ contributions

AIG and SMM designed and implemented SprayNPray. CRA, VSC, SEL, and JMB generated data for case studies. AIG, CRA, and SMM wrote the early version of the manuscript. AIG, CRA, VSC, SEL, JMB, and SMM read, commented, edited, and approved the final manuscript.

## Acknowledgements

We thank Gustavo Ramirez and Michael Pavia for providing many comments, ideas, and insights on this software and manuscript. We thank John McCutcheon for valuable insights into identification of endosymbionts and horizontal gene transfers in eukaryotic assemblies. We also thank Ben Tully for providing comments and insights on the Bin Validation section of this software and manuscript. This is JISAO contribution no. 2021-1139 and PMEL contribution no. 5246.

---

**Supplemental Figure 1:**
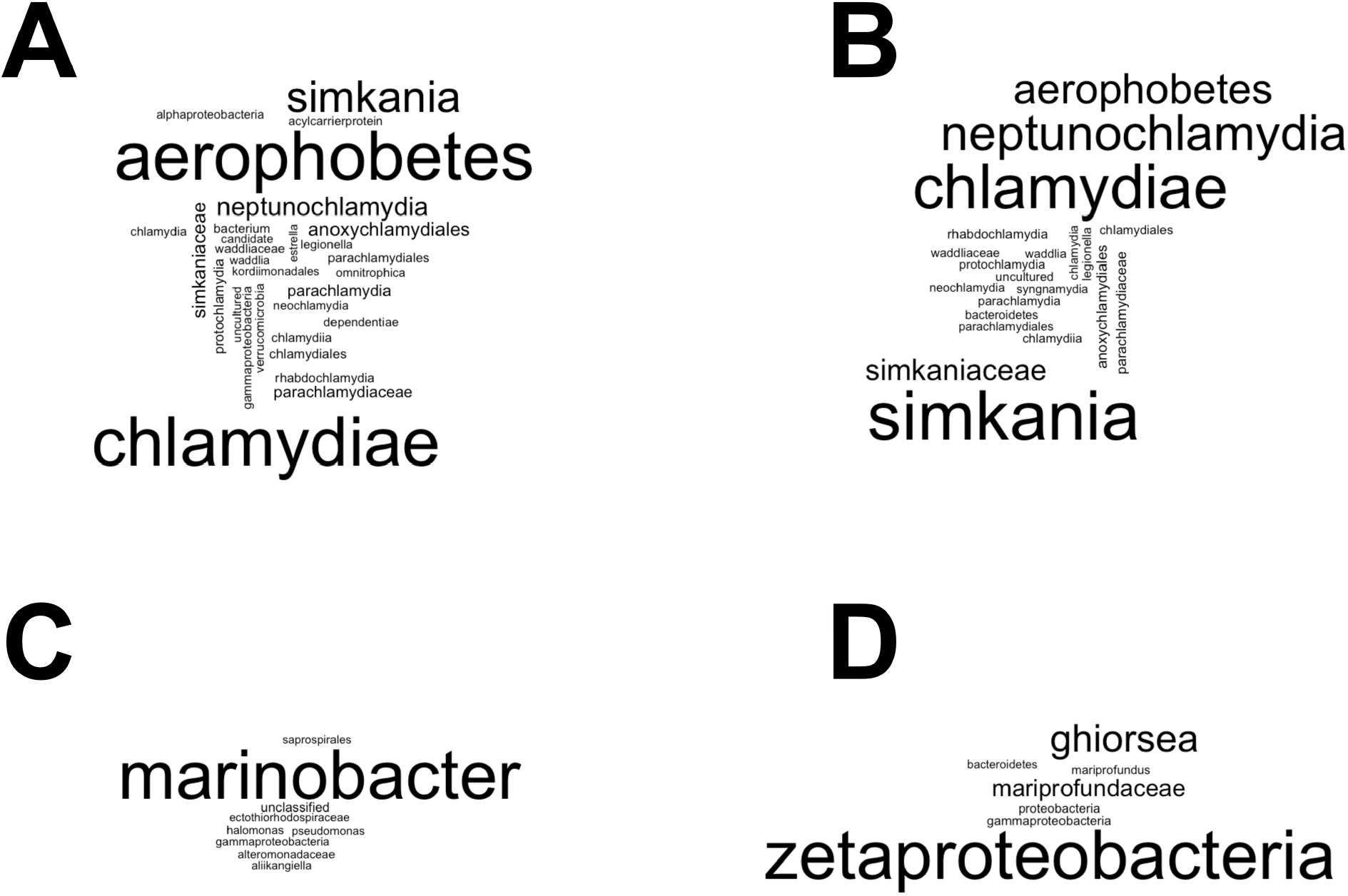
Word clouds generated with SprayNPray based on analysis of four of the MAGs from North Pond described in section 3.1’s case study (Tully et al., 2018): A) NORP81, B) NORP 91, C) NORP151, D) NORP 148. This graphic is generated using the R package “wordcloud” (Fellows, 2018, https://cran.r-project.org/web/packages/wordcloud/wordcloud.pdf).

